# Effect of Ionic Strength on the Assembly of Simian Vacuolating Virus Capsid Protein Around Poly(Styrene Sulfonate)

**DOI:** 10.1101/2022.02.25.481942

**Authors:** Roi Asor, Surendra W. Singaram, Yael Levi-Kalisman, Michael F. Hagan, Uri Raviv

## Abstract

Virus-like particles (VLPs) are noninfectious nanocapsules that can be used for drug delivery or vaccine applications. VLPs can be assembled from virus capsid proteins around a condensing agent like RNA, DNA, or a charged polymer. Electrostatic interactions play an important role in the assembly reaction. VLPs assemble from many copies of capsid protein, with combinatorial intermediates, and therefore the mechanism of the reaction is poorly understood. In this paper, we determined the effect of ionic strength on the assembly of Simian Vacuolating Virus 40 (SV40)-like particles. We mixed poly(styrene sulfonate) with SV40 capsid protein pentamers at different ionic strengths. We then characterized the assembly product by solution small-angle X-ray scattering (SAXS) and cryo-TEM. To analyze the data, we performed Brownian dynamics simulations using a coarse-grained model that revealed incomplete, asymmetric VLP structures that were consistent with the experimental data. We found that close to physiological ionic strength, T=1 VLPs coexisted with VP1 pentamers. At lower or higher ionic strengths, incomplete particles coexisted with pentamers and T=1 particles. Including the simulation, structures were essential to explaining the SAXS data.

## Introduction

About half of known virus families have icosahedral capsids which can, under the right conditions, spontaneously self-assemble around RNA or DNA molecules. Virus assembly is an important step in the life cycle of viruses and has become a target for antiviral research^1–13^ and a model for self-assembly in nanotechnology. ^14–23^ Yet many details of this reaction are unknown. Even a simple virus may have about a hundred subunits, combinatorially more intermediates, and many more assembly pathways.

A complete description of virus assembly should include the structures of intermediates, paths between intermediates, rate constants for each step, and the stability of the species. Resolving the empty capsid assembly mechanism is already a very challenging problem,^24–33^ despite recent progress with experimental techniques such as resistive pulse sensing,^34,35^ mass spectrometry,^36^ charge detection mass spectrometry,^37–40^ and solution small-angle X-ray scattering (SAXS).^41–47^ Our lab, however, developed a robust and transparent mechanism for isolating the most probable intermediates at equilibrium and on the assembly path of an empty Hepatitis B capsid.^48^ Using MC simulations, we created a library of representative intermediates and showed that at equilibrium a very limited number of relevant assembly reaction products were selected based on their Boltzmann weights, computed from the self-association standard free energy between subunits.^49^ Our analysis method assumed that intermediates can only be fragments of capsids. This approach, however, is insufficient to describe the assembly of a virus capsid around a template (e.g., a polymer chain) because the number possible intermediates is considerably larger. The assembly reaction involves electrostatic interactions, and hence depends on ionic strength.

In this paper, we determined the effect of ionic strength on the assembly of virus-like particles (VLPs). We assembled simian vacuolating virus 40 (SV40)-like particles by mixing poly(styrene sulfonate) (PSS) with SV40 capsid virus protein 1 (VP1) pentamers. The structure of the assembly products were characterized by solution small-angle X-ray scattering (SAXS) and transmission electron microscopy at cryogenic temperatures (cryo-TEM). In an earlier study,^50^ we showed that simulations may capture experimental trends of virus capsid assembly reactions. To better analyze our experimental data, we performed Brownian dynamics simulations using a coarse-grained model, accounting for the interactions between the capsid protein subunits and between the capsid protein subunits and the polymer template. The simulations provided a library of stable and thus probable intermediates, obtained based on a physical model that was previously shown to match observations on virus assembly around RNA and other polyanions.^51,52^ By docking the atomic model of the protein pentamer subunit and the PSS monomer into the simulated coarse-grained models, we created atomic models of intermediates. Using our analysis software [D+](https://scholars.huji.ac.il/uriraviv/software/d-software),^48,53–56^ we computed the solution scattering curves of each of the atomic models. We then applied optimization algorithms to select the models that best explained the data and to compute their mass fractions, while ensuring the conservation of total mass.

We found that the mass fraction of pentamers that assembled into T=1 SV40 VLPs was maximal near physiological conditions. Decreasing or increasing the solution ionic strength, which changes the interactions between capsid protein subunits and between the capsid protein subunits and the polymer chain, led to a decrease in the fraction of T=1 SV40 VLPs and an increase in the fraction of free pentamers. In addition, the fraction of incomplete asymmetric VLP structures, obtained by the simulations and observed by cryo-TEM, increased. The approach of combining coarse-grained simulations with analysis of SAXS profiles to identify highly asymmetric assembly structures is general, and can be applied to diverse assembly systems.

## Results and Discussion

As a control, we first determined the effect of ionic strength on the assembly reaction of VP1 pentamers (VP_5_) in the absence of PSS. We measured the solution SAXS intensity as a function of the magnitude of the scattering vector, *q*, from solutions of VP1 pentamers at increasing ionic strength (Figure 1a). Above *q* ~ 0.4 the SAXS curve is adequately fit by the computed scattering curve from the atomic model of a VP1 pentamer (Figure S1). At lower *q* values, the intensity became stronger with decreasing ionic strength. The scattering intensity when *q* → 0 is proportional to the concentration of the scattering particles and the average of their molecular weight squared (assuming there is no interaction between the particles). Analysis of the intensity at the lowest scattering angles (Figure 1b) revealed that the average molecular weight of the assemblies decreased with increasing ionic strength. At the highest ionic strength (562mM), the average molecular weight was about two VP1 pentamers.

**Figure 1:**
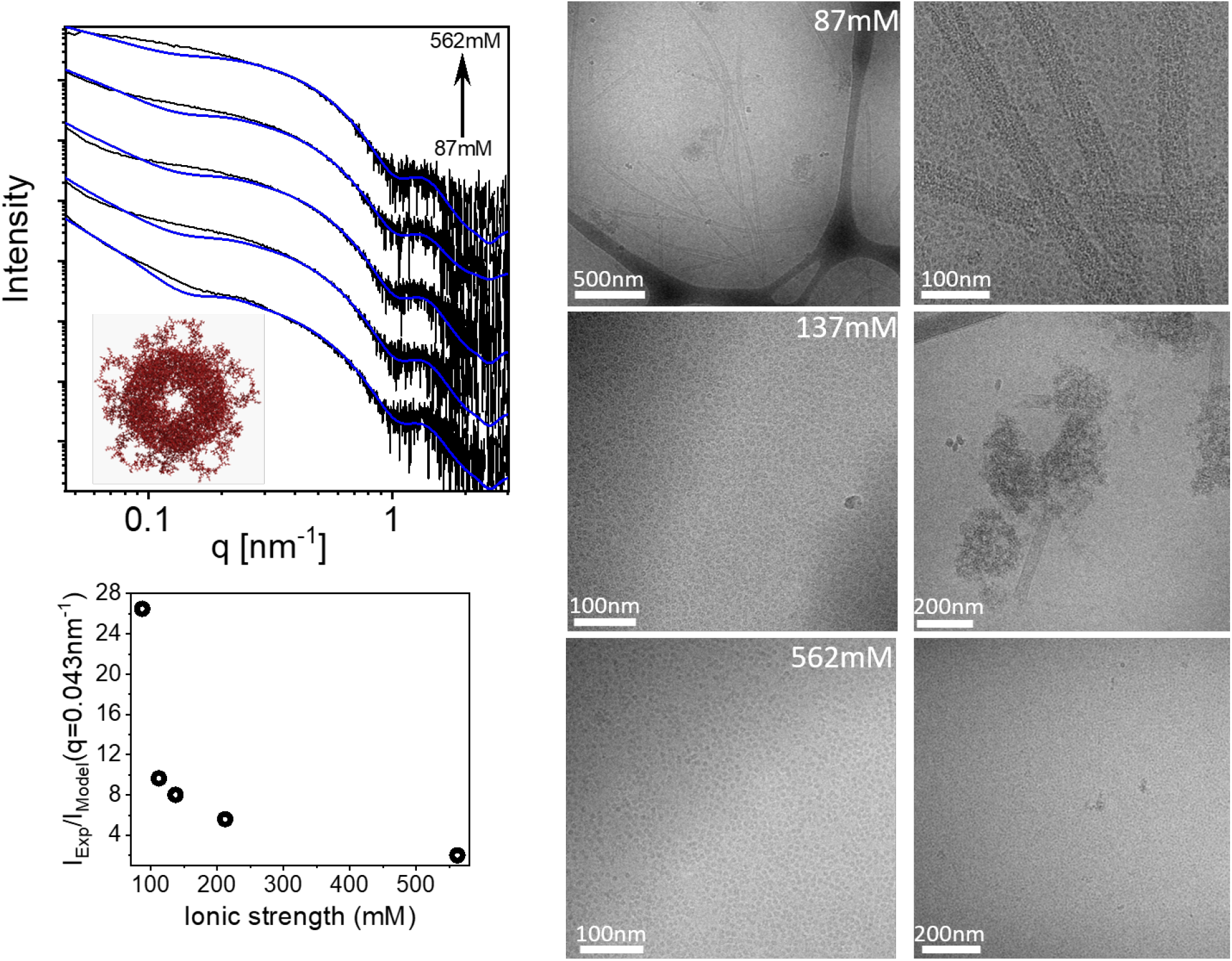
Effect of ionic strength on the assembly of VP1 pentamers, in the absence of poly(styrene sulfonate) (PSS). (a) Small-angle X-ray scattering (SAXS) intensity as a function of the magnitude of the scattering vector, *q*, from 3.75 *μ*M VP1 pentamers at pH 7.2 and varying ionic strengths (black curves). The computed scattering intensity curves (blue) are linear combinations of a *q*^−4^ power law, representing large aggregates, a model of infinitely long tubules with uniform density (inner radius of 15 nm and a wall thickness of 5 nm),^57,58^ and the computed scattering intensity of a soluble VP1 pentamer, based on its atomic structure (see cartoon and red curves in Figure S1).^53,54^ The deviation from the model of soluble pentamer, and as a result, the fraction of tubules and aggregates decreased with ionic strength, in agreement with the intensity at *q* → 0 and cryo-TEM images. (b) The ratio at *q* = 0.043 nm^−1^ between the experimental SAXS signal (*I*_Exp_) and the modeled scattering intensity from soluble VP1 pentamer (*I*_Model_, red curves in Figure S1) as a function of ionic strength. At *q* → 0, the ratio 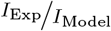 provides a qualitative measure for the average mass of the particles in the sample. (c) Cryo-TEM measurements of 6 *μ*M VP1 pentamers at ionic strengths of 87, 137, and 562mM (a pair of images for each ionic strength condition).

Cryo-TEM images at low salt concentration revealed long tubular and aggregated structures (Figure 1c), which contribute to the enhanced and slightly oscillatory scattering intensity at low *q* (Figure 1a). The measured scattering intensities could be modeled as a coexistence of soluble VP1 pentamers, VP1 tubules, and VP1 aggregates (Figure 1a, blue curves). The mass fraction of soluble VP1 pentamers decreased by about 20% when decreasing the ionic strength from 562 to 87mM.

We then mixed PSS with VP1 pentamers and measured the SAXS curves as a function of ionic strength (Figure 2a). The scattering curves are consistent with the formation of T=1 SV40 virus-like particles (VLPs).^47,50^ Comparing the scattering curves (Figure 2b) shows that at low ionic strength, the scattering intensity at low angles is stronger than at higher ionic strength. This result is consistent with a higher average molecular weight at lower ionic strength, as observed with VP1 pentamers in the absence of PSS (Figure 1a). In addition to T=1 VLPs, excess free VP1 pentamers contributed to the scattering intensity. Nevertheless, a linear combination of T=1 VLPs and free VP1 was insufficient to explain the data, indicating that other structures coexisted. The number of possible structures that may form when PSS interacts with VP1 pentamers is immense because of the number of possible PSS chain conformations and the number of ways that each conformation can interact with the VP1 pentamers. At equilibrium, however,the number of dominant structures is likely to be much smaller, and their mole fraction is set by their excess stability with respect to the reactants (PSS and pentamers). ^50^

**Figure 2:**
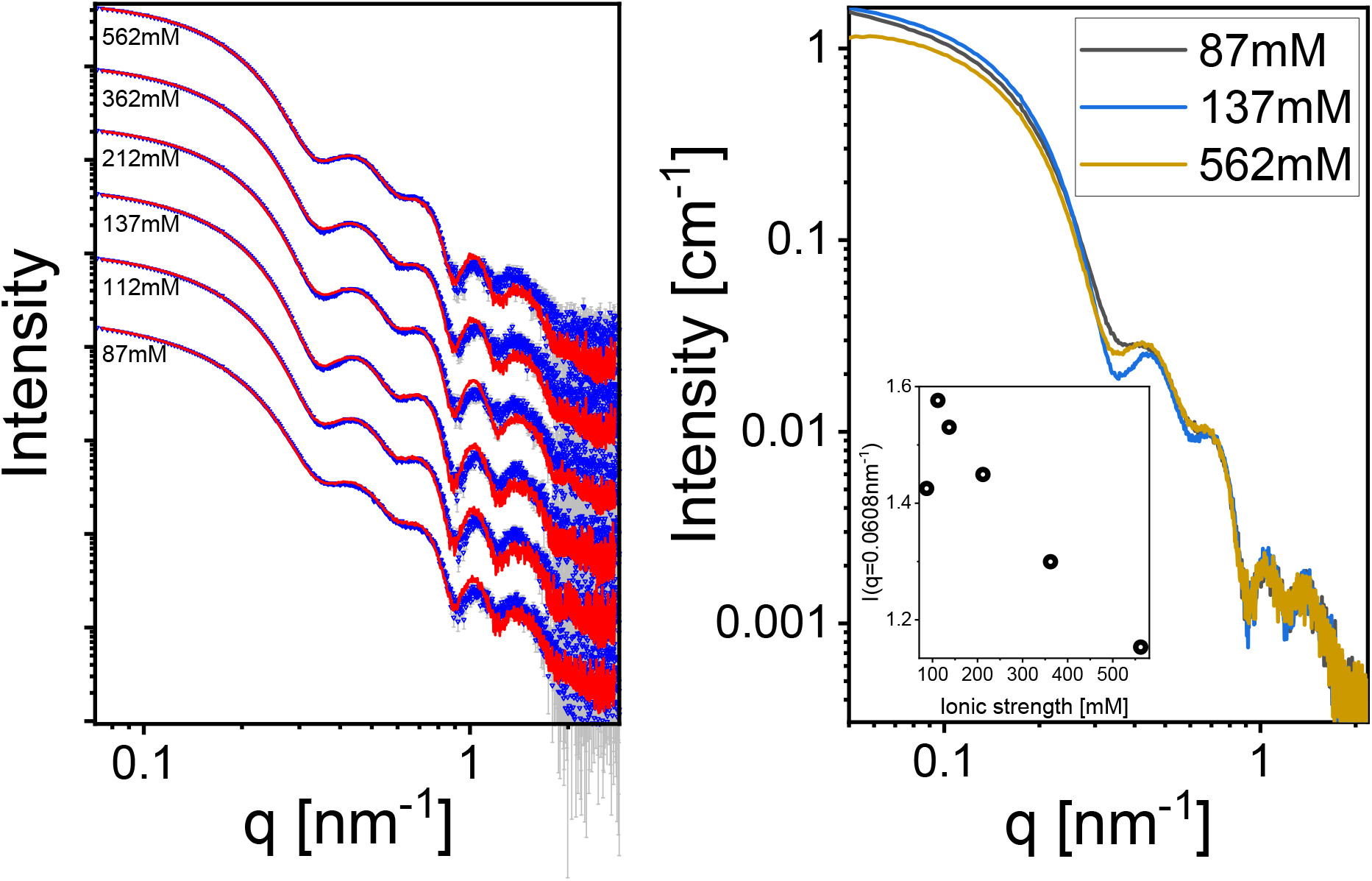
Assembly of VP1 pentamers and PSS at different ionic strengths. Assembly was induced by mixing equal volumes of 0.87 *μ*M 78kD PSS and 13 *μ*M VP1 pentamer aqueous solutions at different ionic strength conditions (see Materials and Methods). Each SAXS curve was fit to a linear combination of the following SAXS curves: Scattering models of solvated PSS-containing T=1 VLP and of free VP1 pentamer, both calculated using [D+](https://scholars.huji.ac.il/uriraviv/software/d-software) software;^53,54^ normalized scattering signal of VP1 pentamers at the corresponding ionic strengths; and simulated models, representing possible assembly products (for a complete explanation of the pool of models see Materials and Methods). (b) Comparing the SAXS curves from representative assembly conditions, showing the nonmonotonic changes in the measured scattering curves as the ionic strength of the solution was increased. The inset shows the absolute intensity at a low scattering angle (*q* = 0.0608 nm^−1^) as a function of ionic strength, corresponding to the nonmonotonic change in the particles’ average mass.

To rationally unravel the dominant coexisting stable structures and analyze our scattering data (Figure 2), we performed Brownian dynamics simulations using a coarse-grained model for templated assembly of VP1 pentamers around a PSS chain. The simulation model for pentamer-pentamer interactions accounts for excluded volume and short-ranged attractions that drive assembly toward T=1 capsids. The pentamer-PSS and PSS-PSS interactions include excluded volume and screened electrostatics (see Materials and Methods for details). We performed the simulations under different ionic strengths and pentamer-pentamer attraction strengths.

When the simulations attained steady-state, we isolated the assembled VLPs (Figure 3a) and classified them according to their size (determined by the number of VP1 pentamers attached to the PSS chain).

**Figure 3:**
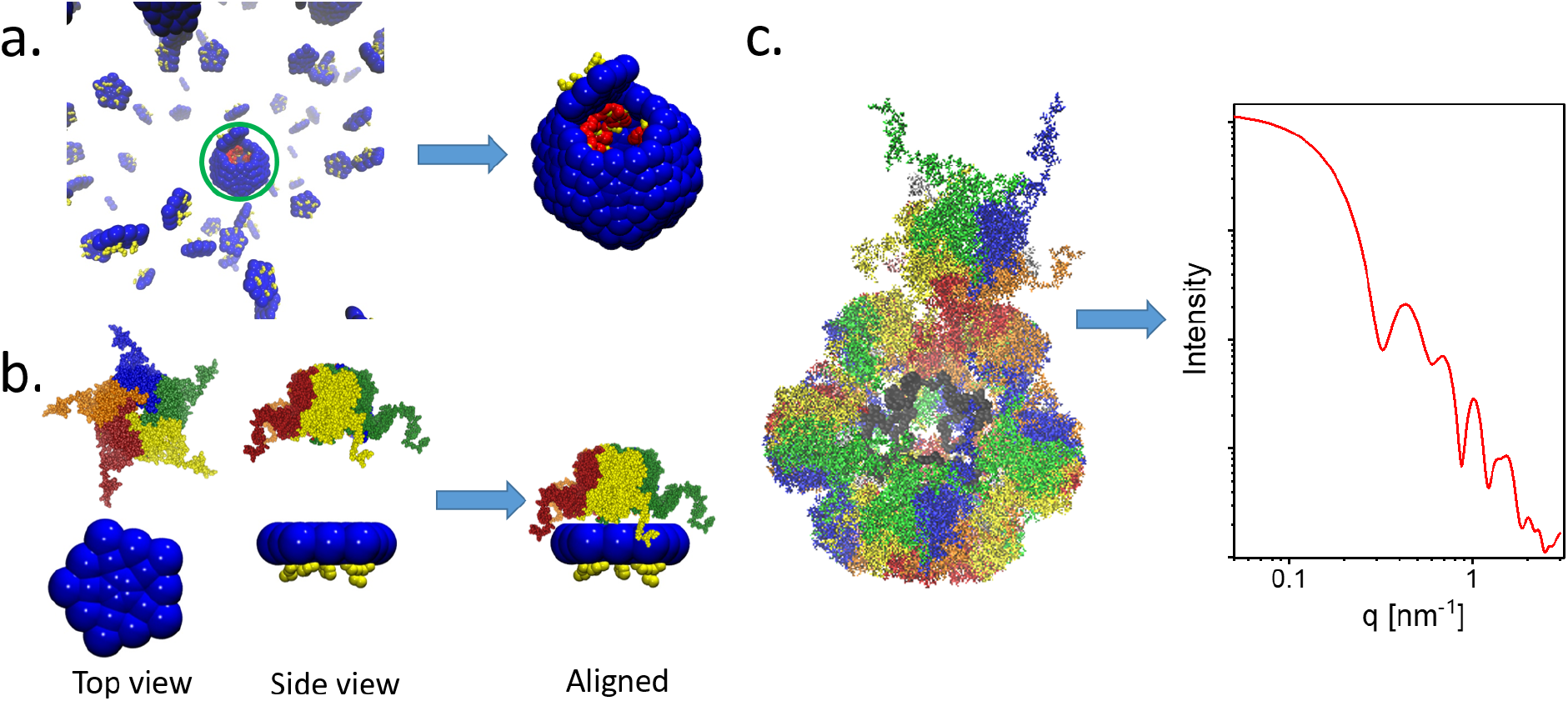
Illustration of our protocol for converting the simulated coarse-grained models to atomic models for computing their solution X-ray scattering curves. (a) Isolating the assembled particle (in the blue circle) from the simulation box after steady-state was attained. (b) Scaling the simulation pentamer coarse-grained model to the real-space atomic structure of VP1 pentamer (PDB ID 1SVA) and aligning the atomic structure to a reference coarse-grained subunit. (c) Computing the solution X-ray scattering curve from an atomic model of a VLP particle. The aligned reference atomic VP1 pentamer model (b) was docked into the isolated coarse-grained VLP assembly (a), accounting for symmetry by defining the rotations and translations of repeating pentamer subunits. The solution scattering curve of the VLP atomic model was computed by [D+](https://scholars.huji.ac.il/uriraviv/software/d-software) software.^53,54^ The hydration layer of the subunits and the instrument resolution function were taken into account as described in Refs.^48,55,56^

We docked the atomic model of a VP1 pentamers (Figure 1a) into the simulated coarse-grained model pentamer (Figure 3b), after scaling the simulations to the correct physical dimensions (according to the atomic model of the subunit). Similarly, we docked the atomic model of a styrene sulfonate monomer into the simulated coarse-grained PSS chain model.

We then aligned the atomic models onto each of the isolated coarse-grained simulated VLP subunits (Figure 3a). We thereby created an atomic model of the simulated VLP particle and calculated its expected solution X-ray scattering curve, using our home-developed [D+](https://scholars.huji.ac.il/uriraviv/software/d-software) software^53,54^ (Figure 3c).

The morphologies of the simulated VLP assemblies varied with the strength of the interaction between pentamers and ionic strength (Figure 4). We ran a large number of simulations under a wide range of conditions and created a diverse and representative library of stable self-assembled structures. The solution X-ray scattering curve from the atomic model of each structure was computed as demonstrated (Figure 3c) and a library of representative scattering curves was created (Figure 5). In addition, we computed a series of scattering curves from models of incomplete empty T=1 VLPs, containing between 1 and 12 VP1 pentamers on the T=1 lattice (Figure S2). The typical T=1 oscillations are observed when the capsid is half complete or larger (Figure S2).

**Figure 4:**
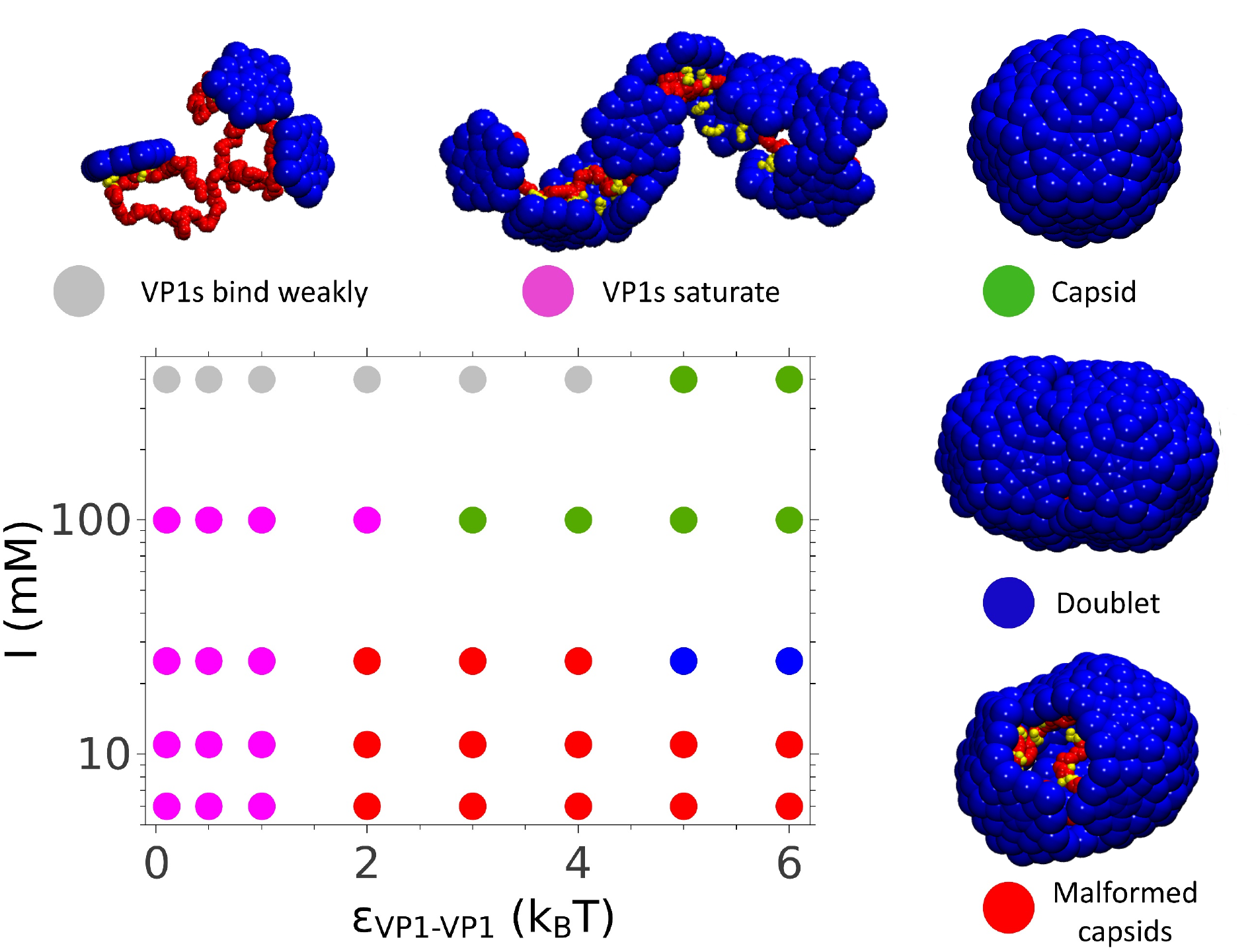
The predominant assembly morphology as a function of VP1-VP1 affinity, *ϵ*_VP1-VP1_, and the ionic strength, *I*, predicted by the coarse-grained simulations. VP1s were modeled as rigid pentamers, interacting with 78 kDa PSS, modeled as a 400-mer bead-spring poly-electrolyte. Morphologies are indicated by symbol color: VP1s bind weakly to the PSS (gray); VP1s saturate the PSS, but do not nucleate (pink); VP1s package the PSS in capsids with T=1 symmetry (green); VP1s package the PSS in two capsids (Doublet), each with T=1 symmetry (blue); Overly strong interactions drive excessive nucleation and formation of malformed capsids (red).

**Figure 5:**
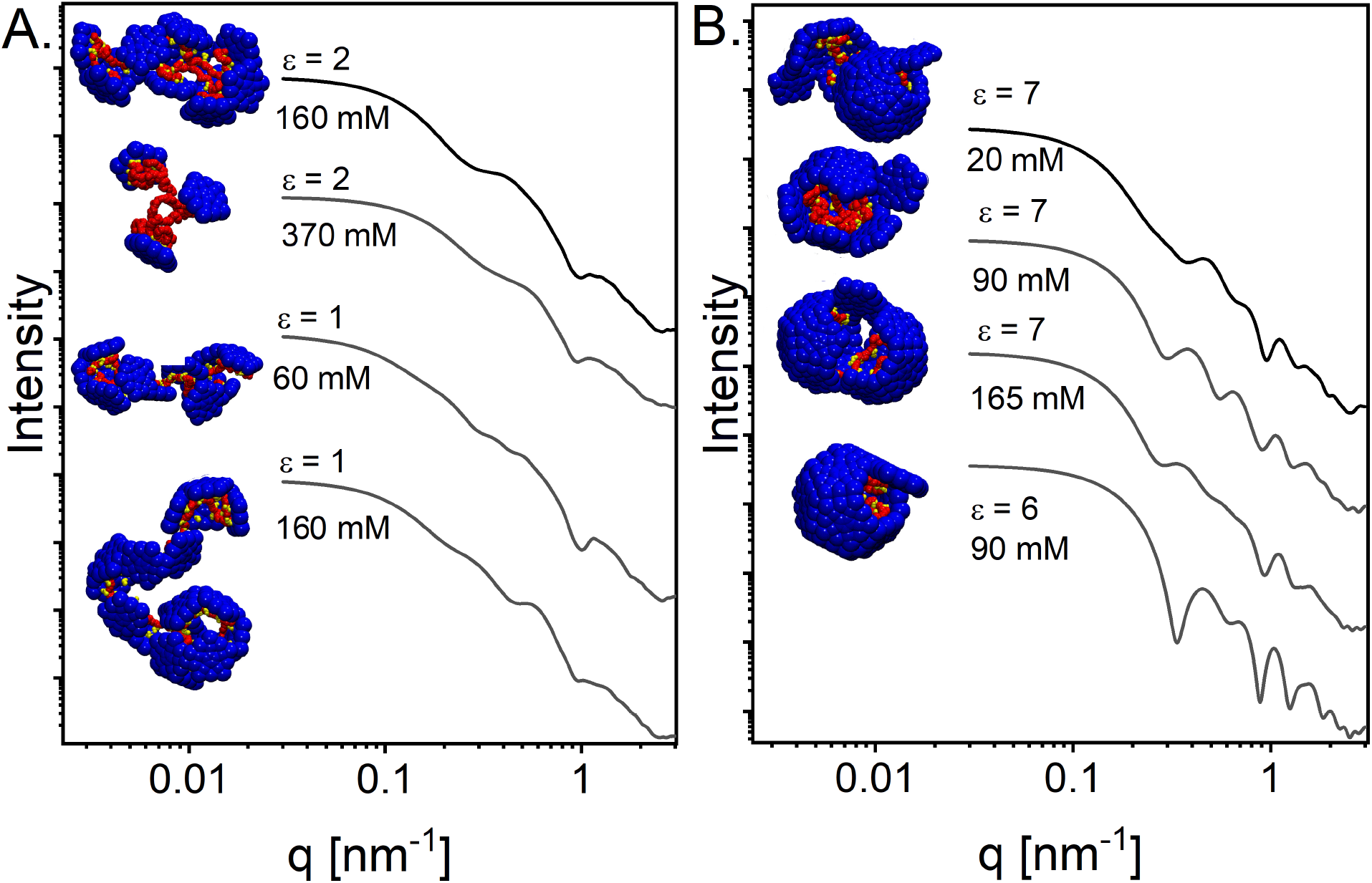
Solution X-ray scattering curves from simulated VLPs under different ionic strengths and interaction strength between pentamers, *ϵ*, as indicated. The scattering curves were computed based on atomic models of the coarse-grained VLPs as demonstrated in Figure 3.

When using the scattering curves of T=1 SV40 VLPs, VP1 pentamers, and the simulated assemblies, we were able to adequately fit the data (Figure 2a). The fraction of T=1, VP1 pentamers, and simulated assemblies varied with ionic strength (Figure 6a,b). The structures of the coexisting simulated assemblies were similar to the particles observed by cryo-TEM (Figure 6c–e). The mass fraction of the simulated assemblies decreased with ionic strength to a minimum at physiological conditions (ionic strength of 137mM) and then increased with further increase of the ionic strength. Even when the mass fraction of the simulated particles was minimal, a linear combination of the computed scattering curves from a complete T=1 SV40 VLP and VP1 pentamers was insufficient to fit the scattering data while maintaining total mass conservation and the correct PSS:VP1 molar ratio (Figure 7). Thus, the simulated assemblies were essential to explain the SAXS profiles.

**Figure 6:**
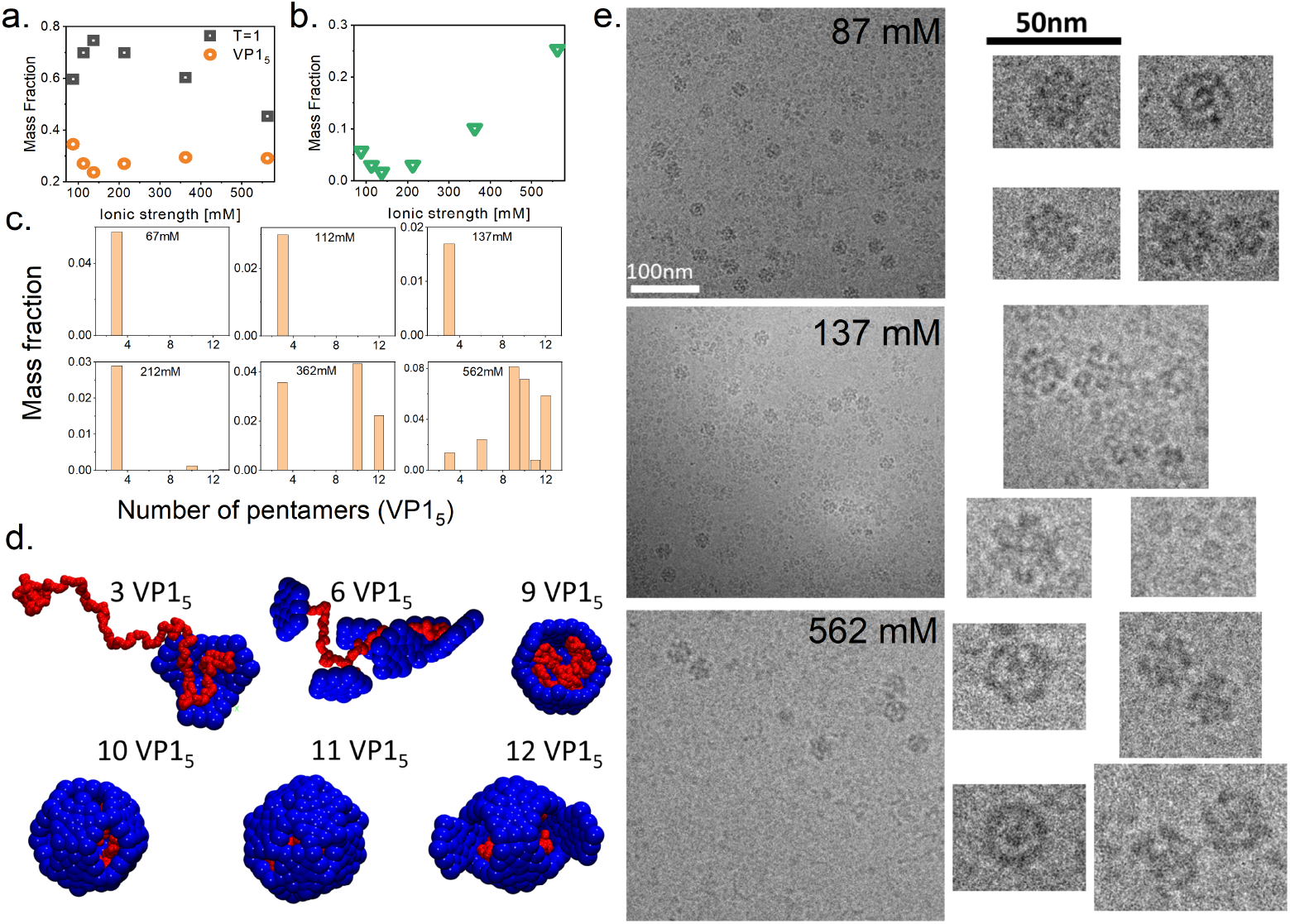
Computed dominant states for assembly of PSS-containing VLPs at pH 7.2 and indicated ionic strengths. (a) Mass fractions of the two dominant populations as a function of ionic strength, based on the SAXS curve fitting procedure. Black symbols correspond to T=1 PSS-containing VLPs and orange symbols correspond to a mixture of free and assembled (Figure 1) VP1 pentamers. (b) Total mass fraction of incomplete simulated PSS-VP1 assemblies as a function of ionic strength. (c) The distributions of the total mass presented in panel b as a function of the size (number of VP1 pentamers) of the simulated assembled states. The ionic strength is indicated in each histogram. (d) The dominant coarse-grained structures that were chosen by the SAXS curve fitting procedure. The number of VP1 pentamers incorporated in each assembled structure is indicated. (e) Cryo-TEM images of assembly reaction products at pH 7.2 and three representative ionic strengths (87, 137, and 562mM). The final concentrations of VP1 pentamers and PSS were 6 *μ*M and 0.4 *μ*M, respectively. For each ionic strength condition (indicated in the Figure) a selected set of particles is shown on an expanded scale.

**Figure 7:**
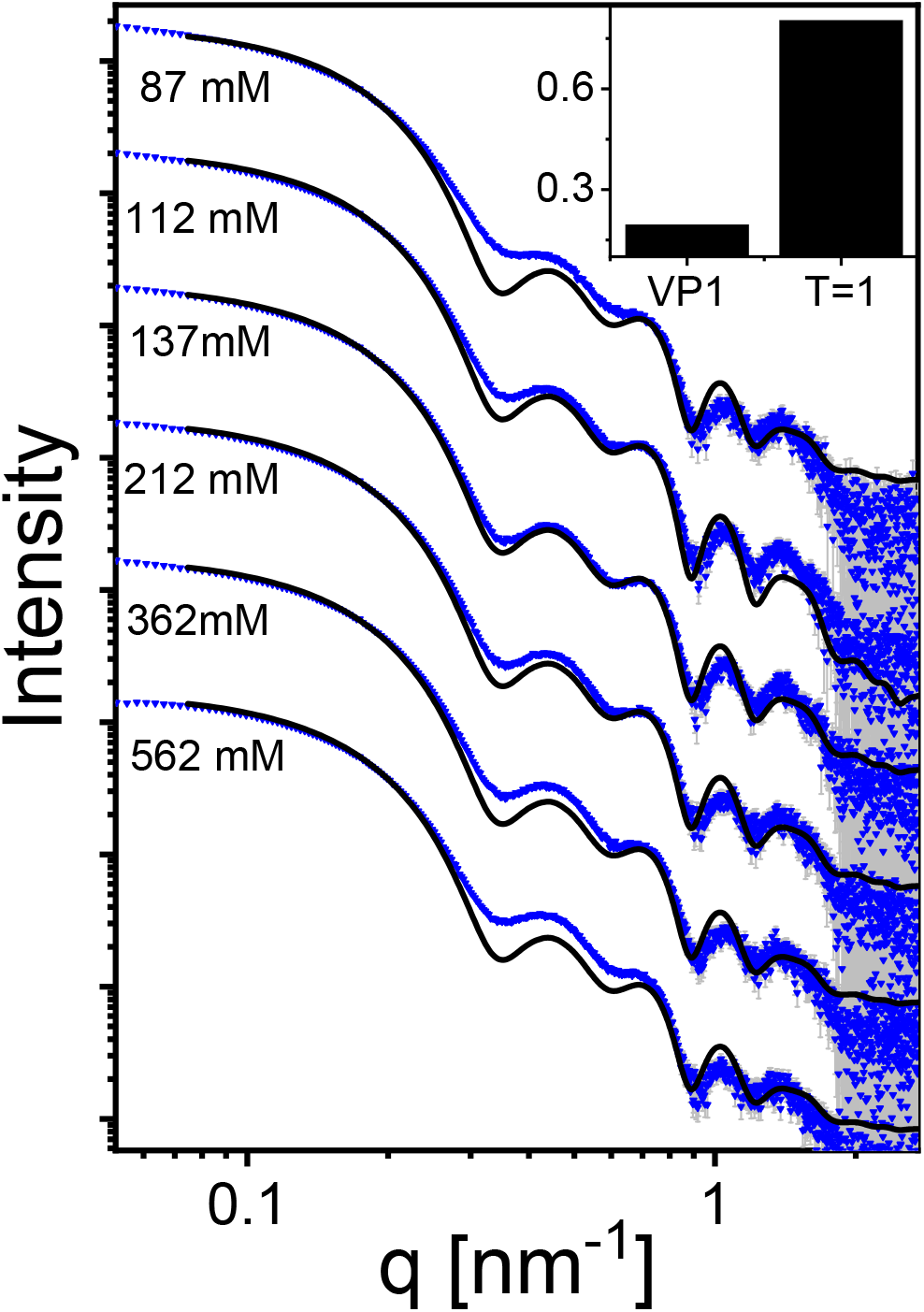
SAXS profiles are poorly fit if simulation assemblies are not included. The plots show fits to the solution X-ray scattering data from Figure 2 using a linear combination of the computed scattering curves from atomic models of VP1 pentamers and complete T=1 SV40 VLPs. Very similar fitting results are obtained when adding to the linear combination the scattering curve from Figure 1 with the corresponding salt solution. The fits were performed while maintaining mass conservation. The inset shows the best fitted mass fraction of the two models, which was very similar in all cases.

At a fixed ionic strength (137mM), we varied the molar ratio between PSS and VP1. Our analysis shows that the contribution of free VP1 pentamers was minimal at PSS: VP1 molar ratio of 1:12 (the expected T=1 stoichiometry) and increased with their molar fraction (Figure 8). The fraction of incomplete simulated complexes, however, was only weakly dependent on the PSS:VP1 molar ratio.

**Figure 8:**
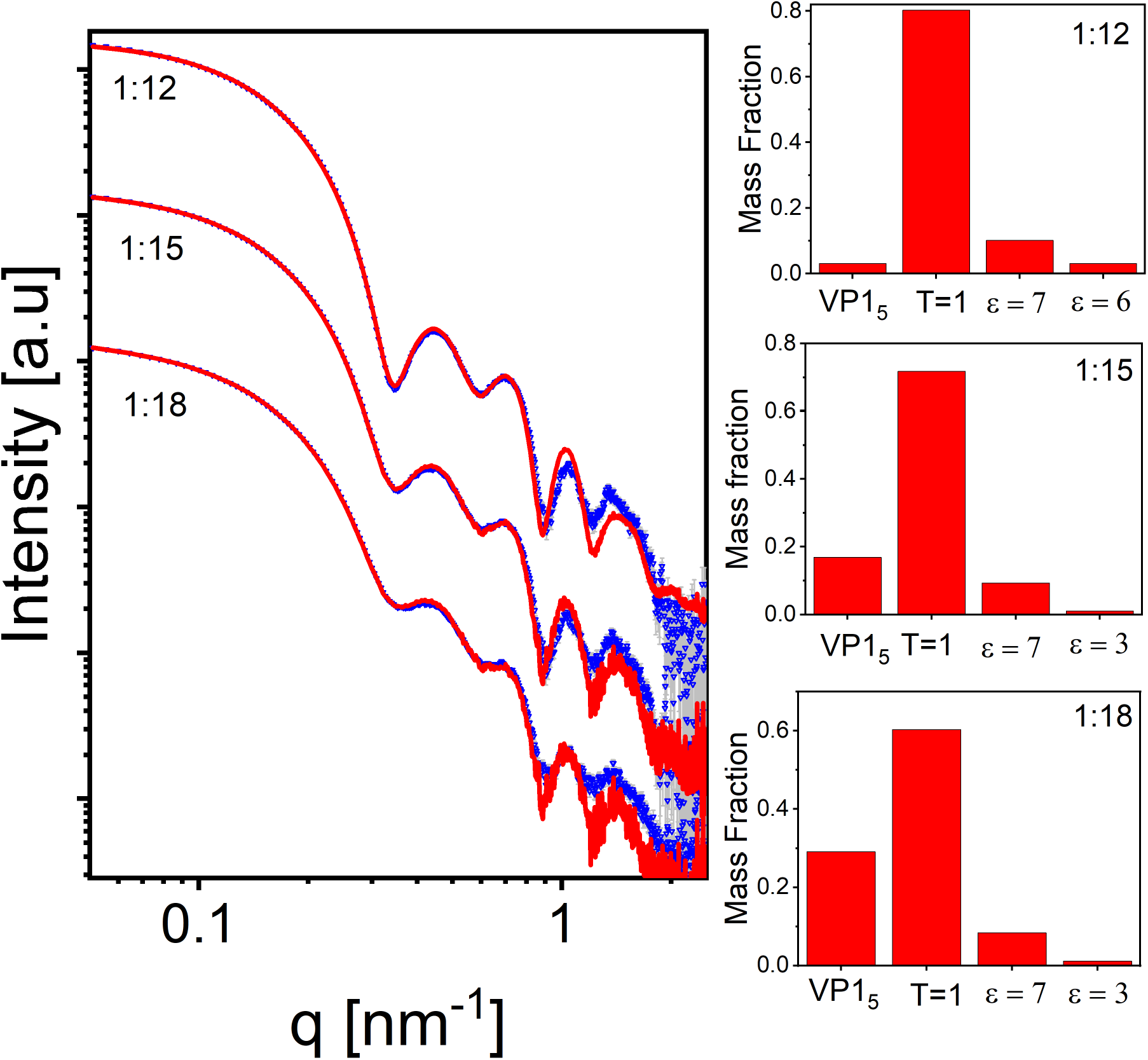
SAXS analysis of assembly reactions at varying PSS to VP1 molar ratios. Assembly was performed at a fixed ionic strength (137mM) and induced by mixing equal volumes of 0.87 *μ*M 78kD PSS solution with VP1 pentamer solution at concentrations leading to the indicated molar ratios. Each SAXS curve was fit to a linear combination of modeled scattering curves of: free VP1 pentamer and the normalized scattering signal of VP1 pentamers at ionic strength of 137mM from Figure 1, solvated PSS-containing T=1 VLP model, and simulated model of VP1 - PSS complexes, representing likely assembly products with the indicated interaction strength, *ϵ*, values.The modeled scattering curves were calculated using [D+](https://scholars.huji.ac.il/uriraviv/software/d-software) software.^53,54^ The histograms show the mass fraction of each model.

## Conclusions

In this paper, we combined SAXS, cryo-TEM, and Brownian dynamics simulations to unravel the effect of ionic strength on the assembly of SV40 VP1 pentamers around a charged polymer template. We found that free VP1 pentamers coexisted with T=1 VLPs as well as incomplete asymmetric structures. Close to physiological ionic strength, the mass fraction of incomplete structures was minimal, consistent with other virus systems (e.g.^27,29,59–65^). Deviation from physiological ionic strength toward lower or higher ionic strengths, which changes the interaction between subunits, increased the mass fraction of incomplete structures. The morphology of the incomplete structures, predicted by the simulations, was consistent with cryo-TEM images and essential for the analyzing the SAXS data. Our approach is broadly applicable to self-assembly reactions, and provides a means to elucidate the ensemble of structures for complex systems involving a large number of subunits and an immense number of possible products. Finally, while we focused on the equilibrium product distribution in this article, the approach can also be applied to time-resolved SAXS to identify the structures of assembly intermediates and their dynamics.

## Materials and Methods

### Sample Preparation

#### VP1 production and purification

VP1 virus-like particles (VLPs) were produced as previously described in Spodoptera frugiperda (sf9) cells, using bacolovirus expression vector. ^66^ VP1-VLPs were then purified from nuclear extract using CsCl density gradient centrifugation. Nuclear extracts were 2-fold diluted with 0.5 M NaCl solution and added to a concentrated CsCl solution, adjusted to a final density of 1.3 g/mL. The nuclear extract suspension was centrifuged in a Beckman 13.2 mL, open-top Thinwall polypropylene tubes (cat. number 331372) in SW41 Ti Swinging Bucket rotor at 38,000 RPM for 40 h. Following centrifugation, VP1-VLPs could be detected by scattered light, as a thick white band at the middle of the tube. The band was pulled and rebanded using a second centrifugation. The fractions were analyzed to asses VP1 content using SDS-polyacrylamide gel electrophoresis (Novex WedgeWell 4-12 % Tris-Glycine) with Coomassie-Blue staining (Instant blue stain, Expedeon). VP1-VLPs were dialyzed against M NaCl at 4 C° using GeBa dialysis tubes with a cutoff of 8 kDa (Gene Bio Application Ltd. cat. no. D070-6). Two dialysis cycles were applied, 1.5 h each, against bulk solutions, whose volumes were 1000X the VP1-VLPs suspension volume. VP1-VLPs were stored up to a month at 4 C°.

#### Preparing VP1 pentamers solution for assembly reactions

The assembly of polymer-containing VLPs (pcVLPs) followed the disassembly-reassembly procedure as described earlier.^47^ Briefly, the purified VP1-VLPs were first disassembled, using two dialysis cycles against disassembly buffers. VP1-VLPs were first dialyzed against disassembly buffer A, containing: 20mM Tris at pH 8.9, 2mM DTT, 5mM EDTA and 50mM NaCl, followed by a second dialysis against disassembly buffer B, which was similar except for a lower concentration of EDTA (2mM instead of 5mM). Both dialysis cycles were performed at 4 C° for 1.5-2 h each, where the ratio between the bulk solution volume and the VP1-VLPs suspension volume was greater than 1000. Following disassembly, the solution containing VP1 pentamers was centrifuged at 20,000 g for 40 min at 4 C° to precipitate larger oligomers. The concentration of VP1 pentamers was measured using UV-Vis absorption spectroscopy with an extinction coefficient of 32, 890 M^−1^cm^−1^ for a VP1 monomer.

#### VP1-PSS measurements

The reactions were initiated by mixing equal volumes of 13 *μ*M VP1 pentamers in disassembly buffer B (20mM 2-Amino-2-hydroxymethyl-propane-1,3-diol (Tris), pH 8.9, 2mM DTT, 2mM EDTA and 50mM NaCl) with 0.87 *μ*M 78 kDa polystyrene sulfonate (PSS) solutions, containing 100mM 3-morpholinopropane-1-sulfonic acid (MOPS) buffer (pH 7.2) and different concentrations of NaCl. SAXS measurements were performed following incubation for at least 5 h at ambient room temperature.

#### VP1 pentamer assembly in the absence of PSS

PSS-free assembly reactions of VP1 at each solution condition were initiated by mixing equal volumes of 7.5 *μ*M VP1 pentamers in disassembly buffer B with buffered solutions, containing 100mM MOPS buffer at pH 7.2 and different NaCl concentrations. All samples were measured following several hours of incubation at ambient room temperature.

### SAXS Measurements

Solution small-angle X-ray scattering (SAXS) measurements were performed at the P12 EMBL Beamline (headed by D. Svergun) in PETRA III (DESY, Hamburg).^67^ Measurements were taken using an automated sample changer setup as described.^48,49,68^ The wavelength of the incident X-ray beam was 1.24 Å and the scattering intensity was recorded on a single-photon PILATUS 2M pixel area detector (DECTRIS). The sample to detector distance was 3.1 m. 30-40 *μ*L of each sample were injected in each measurements and 25 frames were recorded with an exposure time of 45 ms per frame. Initial reduction of the scattering signals to one dimensional curves of scattering intensity as a function of the magnitude of the scattering vector, *q*, was performed using the P12 pipeline.^69^

Background measurements before and after each sample were performed on the solvent of each sample, under identical measurement conditions. Both background and sample scattering curves were averaged over all the frames and the averaged background signal was subtracted from the averaged sample, and gave the final background subtracted scattering intensity curve of the assembly reactions, as explained in our earlier papers.^48,53–56^ All the assembly reactions were measured at 25 °C.

SAXS Measurements were also performed at the ID02 beamline (headed by T. Narayanan) in the European synchrotron radiation facility (ESRF, Grenoble). Measurements were taken using the flow-cell setup, which included a temperature controlled, 2mm thick, quartz capillary.^70,71^ The wavelength of the incident beam was 0.995 Å and the scattered intensity was recorded on a Rayonix MX170-HS detector.^71,72^ Data reduction was performed by SAXSutilities software.

### Cryo-TEM measurements

Assembly reactions for cryo-TEM measurements were prepared using a protocol that is identical to the protocol used for SAXS measurements. The final concentration of VP1 and PSS were 6 *μ*M and 0.4 *μ*M, respectively. All assembly reactions were incubated for ~24 h at ambient room temperature. Samples preparation and image acquisition were performed as described in our earlier publication.^42^

### Coarse-grained computational model

To develop a library of intermediate structures, we performed Brownian dynamics simulations with a coarse-grained model for assembly of T=1 SV40 VLPs around a polyelectrolyte. The model is adapted from models previously used to simulate assembly of empty capsids,^73–75^ which were extended model assembly around RNA and other substrates in Refs.^51,52,76–79^

The T=1 SV40 VLP capsid is modeled as a dodecahedron composed of 12 pentagonal subunits. The subunits are attracted to each other by an attractive Morse potential between Attractor (‘A’) pseudoatoms located at each subunit vertex. The Top (‘T’) pseudoatoms interact with other ‘T’ psuedoatoms through a potential consisting of the repulsive term of the Lennard-Jones (LJ) potential, the radius of which is chosen to favor a subunit-subunit angle consistent with a dodecahedron (116 degrees). The Bottom (‘B’) pseudoatom has a repulsive LJ interaction with ‘T’ pseudoatoms, to prevent ‘upside-down’ assembly. The ‘T’, ‘B’, and ‘A’ pseudoatoms form a rigid body.^73–75^ To represent the capsid shell excluded volume more accurately than the original model, we add a layer of ‘Excluder’ pseudoatoms, having a repulsive LJ interaction with the polyelectrolyte and the ARMs (discussed next). The attraction strength is controlled by the model parameter *ε*_ss_. In Ref.,^52^ the relationship between the potential well-depth *ε*_ss_ and the dimerization standard Helmholz free energy was estimated to be 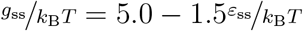.

We represent the PSS with a linear bead-spring polyelectrolyte, with a charge of −*e* per bead. To represent the SV40 capsid protein RNA binding domains (arginine rich motifs, ARMs), we affix flexible polymers to the inner surface of the model capsid subunits. There are 5 ARMs per capsid subunit (since it is a homopentamer), and each ARM contains 5 positively charged beads and 0 charge-neutral segments. The first segment of each ARM is part of the subunit rigid body, whereas the remaining ARM segments are not rigid but connected to the rigid body through the bonded interactions within each ARM. Each ARM is modeled as a bead-spring polymer, with one bead per amino acid. The ‘Excluders’ and first ARM segment are part of the subunit rigid body. ARM beads interact through repulsive LJ interactions and, if charged, electrostatic interactions modelled by a Debye-Hückel (DH) potential.

Electrostatics are modeled using DH interactions, where the Debye screening length (*λ*_D_) is determined by the ionic strength *I_S_* as 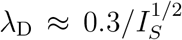 with *λ*_D_ in nm and *I_S_* in molar units. We consider monovalent salt, for which *I_S_* is given by the salt concentration *C*_salt_. Perlmutter et al.^51^ showed that DH interactions compare well to simulations with explicit counterions for the parameter values under consideration; however, we note that the DH approximation is less accurate at lower salt concentrations.

#### Simulations and units

Simulations were performed with the Brownian Dynamics algorithm of HOOMD, which uses the Langevin equation to evolve positions and rigid body orientations in time.^80–82^ Simulations were performed using a set of fundamental units. The fundamental energy unit is selected to be *E*_0_ ≡ *k*_B_*T*. The unit of length *D*_0_ is set to the circumradius of a pentagonal subunit, which is taken to be *D*_0_ ≡ 5 nm so that the dodecahedron inner radius is 1.46*D*_0_ = 7.3 nm is on the order of the size of a SV40 T=1 VLP. Assembly simulations were performed at least 10 times for each set of parameters, with each simulation concluded at either completion, persistent malformation, or 1 × 10^8^ time steps. For all dynamics simulations there were 30 subunits with box size=75 × 75 × 75 nm^3^, resulting in a concentration of 120 *μ*M.

### Fitting SAXS signals

#### Calculation of SAXS intensities from coarse-grained Brownian dynamics simulations

To compute the expected solution X-ray scattering intensity curves from the polymer-VP1 pentamer complex in [D+](https://scholars.huji.ac.il/uriraviv/software/d-software) software,^53,54^ we used the atomic model of a VP1 pentamer (taken from PDB ID 1SVA) and a small sphere representing the PSS monomer. We docked the VP1 pentamer atomic model into its assembly symmetry, describing the manner by which copies of a subunit are shifted and rotated.

#### Generating assembly symmetry files from the simulation frames

An equilibrated MD simulation trajectory included 4 saved frames. Each frame included the size and shape of the box (3 edges and 3 angles between them) and a list of 3D bead coordinates (a polymer contains 400 beads, and the 30 VP1 pentamers each have 22 beads). Each pentamer, contained a list of 22 consecutive bead coordinates (using periodic boundary conditions), 20 of which defined the rigid core whereas the top and bottom beads defined its inner and outer faces, respectively. The coarse-grained representations of the N-terminal ARMs were not included in the simulation frames. The polymer interacted with the pentamers and formed complexes that coexisted with free pentamers.

A Python program identified the pentamers that formed a complex with the polymer, and computed, based on the MD simulation frames, the assembly symmetries of the VP1 pentamers and the polymer subunit. An assembly symmetry contains the manner by which copies of a subunit are shifted and rotated and was required for computing the X-ray scattering curve of the complex in [D+](https://scholars.huji.ac.il/uriraviv/software/d-software) software (Figure 3).

Each equilibrated simulation frame was corrected for periodic boundary conditions by surrounding the simulation box with 26 copies of its translated pentamer beads.

We identified the coordinates of fragmented subunits near the periodic boundaries if the distance between beads and the geometric center of the pentamer was larger than the pentamer enclosing radius. To correct the coordinate list of these subunits, we applied a k-means clustering algorithm with an initial guess of 8 clusters.^83^ If the separations between the resultant centroids were smaller than the size of a pentamer, the clusters were merged, and the number of clusters was modified. A second k-means procedure was applied with the modified number of clusters as an input parameter and the resultant clusters defined the subunit fragments. The fragments were combined by a series of translations.

To isolate the VP1 - polymer complex (Figure 3a), as a first step, VP1 pentamer subunits within an interaction distance of 1.5 in the simulation units (1 in the simulation units equals nm) between the geometric center of the pentamer and the polymer were considered to be part of the complex. The remaining pentamers were initially considered free. A free pentamer, which interacts with any of the pentamers in a complex was added to the complex. This was repeated until no free pentamer had to be added. The interaction cutoff distance between two pentamers was set to 0.6 in the simulation units between the most peripheral beads of these pentamers (the 5 peripheral beads of each pentamer were located at the largest distance from its geometric center).

A reference subunit was aligned such that its geometrical center was at the origin and its director (the vector connecting the bottom and top bead centers) was aligned with the positive *z* axis (Figure 3b). To extract the position and orientation of each pentamer in the complex, the rotation Euler angles and translation vectors were calculated with respect to the aligned reference subunit. The output of the calculation included a list of the geometric centers and Euler angles of each pentamer in the complex and the geometric centers of the polymer subunits. The exported positions were multiplied by an optimized scaling factor of 4.6, representing the ratio between the real-space and simulated VP1 pentamer dimensions (obtained by comparison with scattering data), and translated such that the center of mass of the pentamers is located at the origin.

#### Docking of the VP1 pentamer atomic structure

To compute the solution X-ray scattering curves from VP1 - polymer complex in [D+](https://scholars.huji.ac.il/uriraviv/software/d-software) software,^53,54^ we first aligned an atomic model of a VP1 pentamer, taken from PDB ID 1SVA and a published cryo-TEM data,^46^ with the coarse-grained reference subunit. The center-of-mass of the pentamer atomic model was located at (0,0, 2.6 nm) and the alignment of the pentamer C-arms was compared with the attractor beads in the coarse-grained reference subunit. The alignment was validated by comparing the computed scattering curve of a T=1 particle from the dynamics simulations with the computed scattering curve based on a published cryo-TEM T=1 SV40 virus-like particle structure.^46^ The scattering amplitude of the hydrated VP1 pentamer was computed in [D+](https://scholars.huji.ac.il/uriraviv/software/d-software), using a solvent probe radius of 0.14 nm, a hydration layer thickness of 0.2 nm, and a mean electron density of 364 e/nm^3^

#### Docking of the styrene sulfonate monomer

The orientation-averaged scattering intensity from an atomic model of a styrene sulfonate monomer (CID 75905) was computed and fitted to a sphere model with a radius of 0.23 nm, and a mean electron density of 1208 e/nm^3^. The contribution of the PSS polymer was computed by docking the best fitted sphere model into the assembly symmetry of the 400 polymer subunits, obtained from the dynamics simulation.

### Analysis of SAXS data

The scattering data were fitted to a linear combination of the following computed scattering curves:

1. **Computed free VP1 pentamer:** The solution SAXS intensity from the atomic model of a free VP1 pentamer (taken from PDB ID 1SVA) was computed with a hydration shell, computed by [D+](https://scholars.huji.ac.il/uriraviv/software/d-software) program^53,54^ with a solvent probe radius of 0.14 nm, a hydration layer thickness of 0.2 nm, and a mean electron density of 364 e/nm^3^.
2. **Measured VP1 pentamers:** The measured SAXS curve of the VP1 pentamers at the relevant ionic strength (Figure 1a) was used to take into account the coexisting self-associated VP1 pentamers.
3. **T=1 SV40 Virus-like particle:** The atomic model of the T=1 SV40 virus-like particle was created by docking the VP1 atomic pentamer model and the spherical polymer subunits to a complete T=1 particle taken from a simulation frame (as explained above). The structure was further modified by adding an inner uniform sphere, represented by a hyperbolic tangent model with a radius of 5.5 e/nm^3^, maximum electron density of 378 e/nm^3^, and a slope of 0.8, as explained in ref.^84^ Thermal fluctuations were taken into account by allowing the center of mass of the VP1 pentamers to fluctuate about their position. The fluctuation amplitude was randomly drawn from a uniform distribution between −0.5 and 0.5 nm. The scattering model represents an average of 15 generated T=1 particles.
4. **A library of VP1-polymer complexes:** The library was generated by coarse-grained MD simulations, using a range of pentamer-pentamer interaction parameters (3 ≤ *ϵ*_*pp*_ ≤ 7) and Debye lengths (0.5 nm ≤ *λ*_*D*_ ≤ 1 nm).

#### nonnegative constraint Chi-square fitting

The fitting algorithm included a least-squares fitting of the model to the data, where the contribution of each component, representing its mass fraction, had to be non-negative and obey the constraint of total mass conservation.

## Supporting information

draft_VP1PSS_MainText.pdf

## Acknowledgement

We thank Daniel Harries for very helpful discussions. We acknowledge the European Synchrotron Radiation Facility (ESRF) beamline ID02 (T. Narayanan and his team), the Desy synchrotron at Hamburg, beamline P12 (D. Svergun and his team), and Soleil synchrotron, Swing beamline (J. Perez and his team), for provision of synchrotron radiation facilities and for assistance in using the beamlines. This work was supported by Award Number R01GM108021 from the National Institute Of General Medical Sciences. Computational resources were provided by NSF XSEDE computing resources (XStream, Bridges, and Comet) and the Brandeis HPCC which is partially supported by DMR-2011846. R.A. acknowledges support from the Kaye-Einstein Fellowship Foundation.

## Supporting Information Available

The Supporting Information is available free of charge on the ACS Publications website at DOI:

The Supporting Information includes additional supporting figures and details regarding:

