## Supplementary material for "Effect of Ionic Strength on the Assembly of Simian Vacuolating Virus Capsid Protein Around Poly(Styrene Sulfonate)": draft_VP1PSS_MainText.pdf

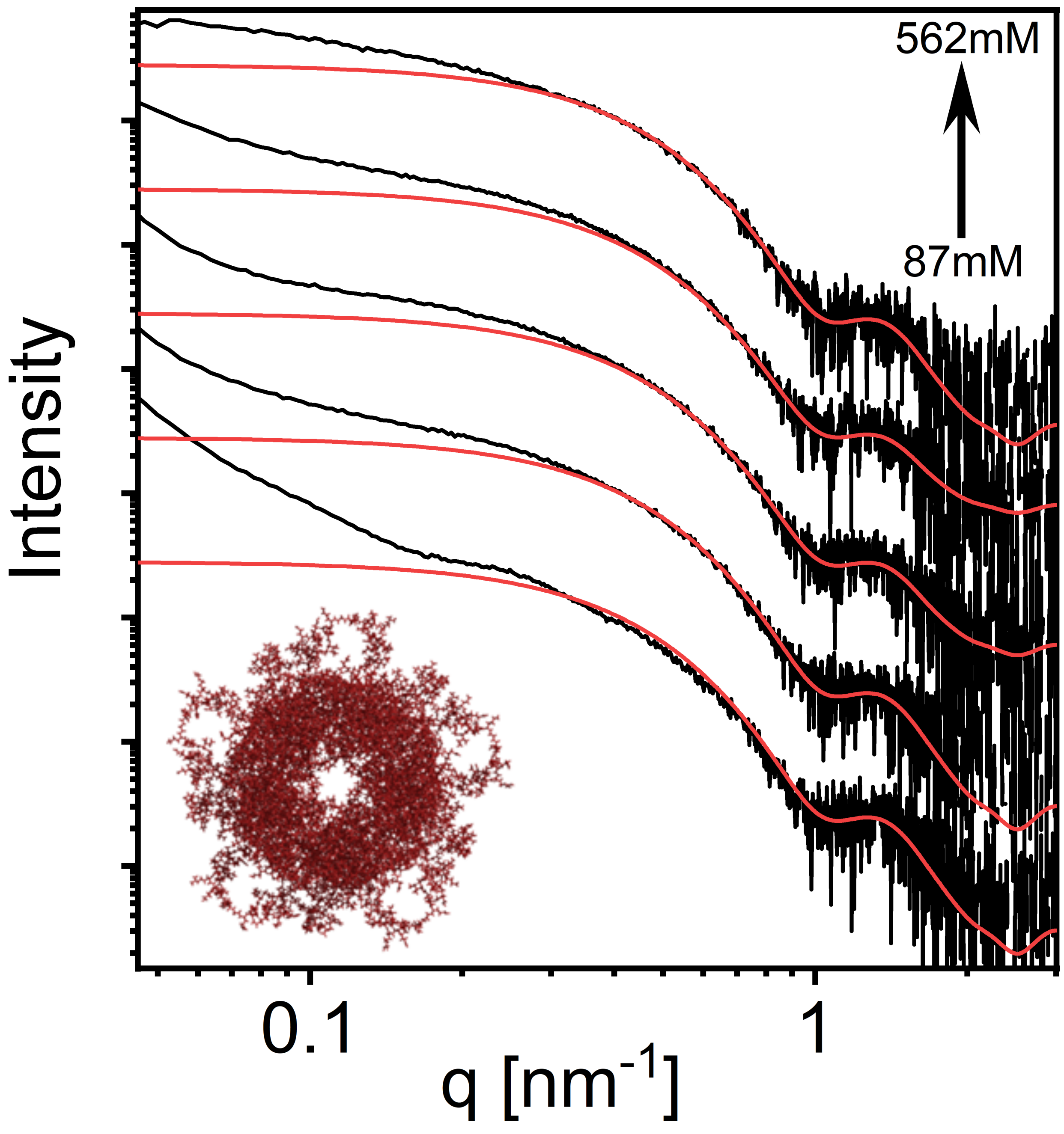


Figure S1: SAXS curves from 3.75 $\mu$M VP1 pentamers at pH 7.2 at varying ionic strengths (black curves). The red curves are the computed scattering curve from a soluble VP1 pentamer, based on its atomic structure, shown at the inset (see Materials and Method section for calculation parameters). The deviations from the red curves at low angles are due to tubular assemblies of VP1 pentamers and VP1 aggregates (Figure 1 of the main text).

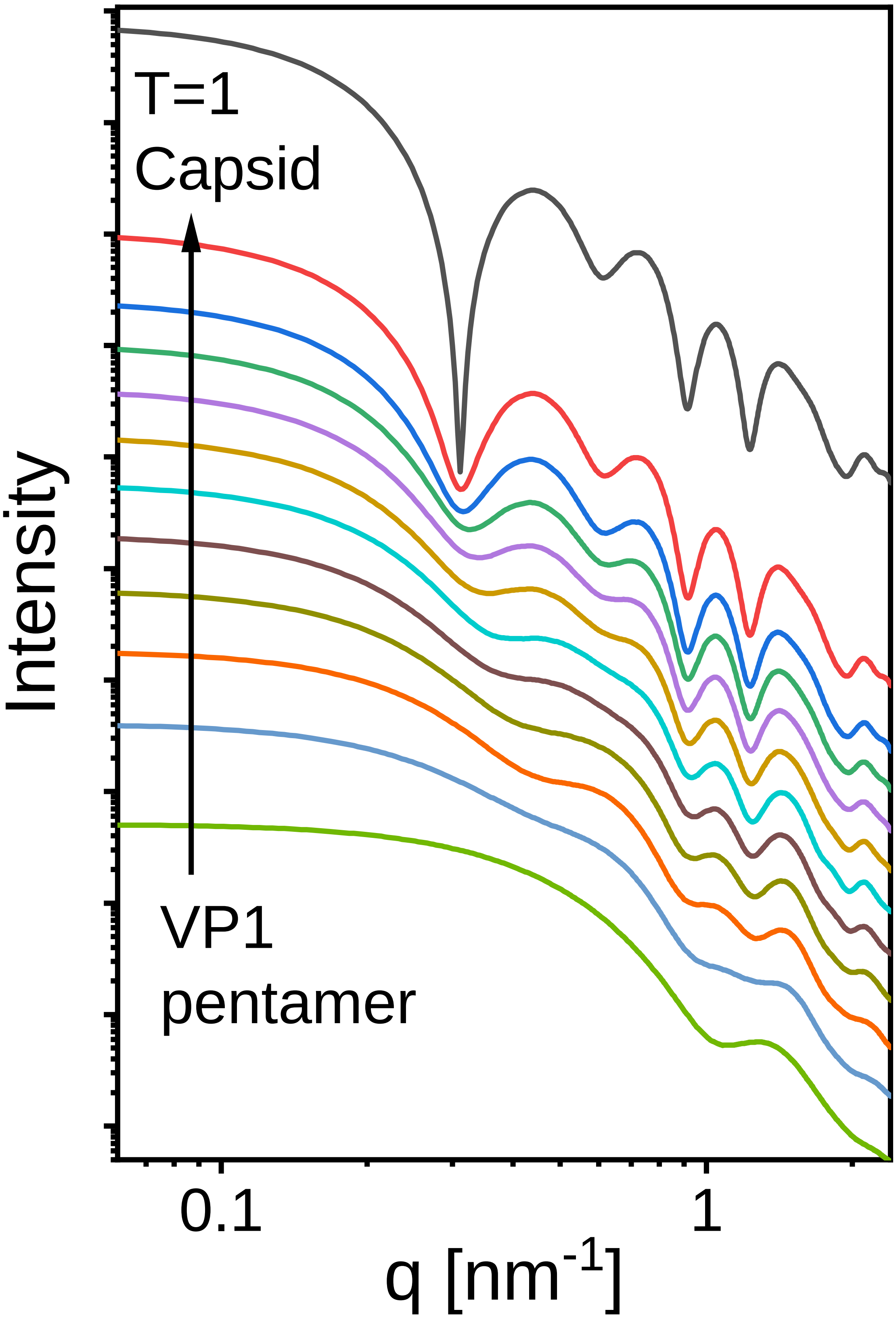




Figure S2: Computed solution X-ray scattering curves from incomplete empty capsids, containing between 1 (bottom curve) and 12 (T=1 capsid, top curve) VP1 pentamers.
